# The Tiling Algorithm – A general method for structural characterization of accurate long DNA sequence reads: application to AAV genome sequences

**DOI:** 10.1101/2025.07.25.666743

**Authors:** Robert E. Bruccoleri, David Rouleau, Celia Slater, Dimpal Lata, Christopher Phillion, Samuel Adjei, Kiran Adhikari, Serena Dollive

**Affiliations:** Congenomics, LLC, Glastonbury, CT, USA; Oxford Biomedica (US) LLC, Bedford, MA, USA

## Abstract

<u>A</u>deno-<u>a</u>ssociated <u>v</u>irus (AAV), a common vector used in human gene therapy, is challenging for DNA sequencing for several reasons. First, AAV’s replication cycle results in structural rearrangements. Specifically, the inverted terminal repeat (ITR) at each end of the genome as well as the genetic payload can invert independently. Second, the ITR can prime DNA replication of the entire viral genome in the gap filling step of a sequencing library preparation protocol. Third, the AAV manufacturing process can produce small fractions of viral particles containing host cell DNA and/or fragments of the helper plasmids. And finally, short read sequencing is ill suited for the characterization of repetitive sequences and structural rearrangements involving several hundred nucleotides. Pacific Biosciences (PacBio) long-read sequencers are capable of full length, accurate, single molecule sequencing of AAV viral genomes, which addresses the challenge of working with repetitive sequences. However, sequence analysis methods based on alignment to a reference are confounded by the other three challenges above. We present a simple algorithm for determining the arrangement of functional elements of single DNA molecules which can be aggregated to provide a sensitive measure of the population of sequences in a sample, including minor species. Using data from four publicly available datasets, we demonstrate our algorithm is able to characterize nearly all of the species in the AAV samples.

## Introduction

<u>A</u>deno-<u>a</u>ssociated <u>v</u>irus (AAV) is a common vector for human gene therapy [1]. Its use in clinical settings requires proof of the purity of any manufactured lot of the virus [2, 3]. Although determination of the DNA sequence via next-generation sequencing (NGS) is not yet required by health authorities as a method for assessing the purity and enumerating contaminants, it is foreseeable that it will be in the future. Certainly, at present, the determination of the DNA sequence is a valuable cross-validation tool along other established assays such as digital droplet PCR in the development of new vectors.

Analyzing AAV genomes using DNA sequencing is difficult for several reasons. First, the virus’s unique replication process [8] rearranges the genetic elements combinatorially. The AAV genome consists of a single stranded DNA molecule with the genetic payload sandwiched between two inverted terminal repeats (ITRs) [9]. The replication process does not conserve the strandedness of the ITRs and the payload resulting in a variety of functional and acceptable vector genomes varying by ITR orientation and strandedness of the expression cassette for each AAV preparation [10]. Second, the ITR’s are close to the read length for short read sequencers such as Illumina’s, and thus, disambiguating the location of a short read containing an ITR can be problematic. Third, the GC content of the ITR as well as common promoters, such as the chicken *beta*-actin promoter, is very high, which limits the use of sequencing methods that rely on PCR amplification. An example of such a promoter is in the self complementary Green Fluorescent Protein (GFP) dataset (folder 2021-scAAV-CBA-eGFP) described in the Material section below.

Sanger sequencing is not appropriate for assessing the broad contents of a manufactured lot because PCR amplification is required, and therefore the Sanger chromatograms represent the macroscopic summation of all the sequences that are amplifiable with a specified set of PCR probes. Thus, Sanger sequencing cannot capture low abundance or non-targeted DNA molecules, including both residual contaminants and sequence variation in the vector [4, 5]. In addition, the read length for Sanger sequencing is too short to capture the entire viral genome, and thus assembly would be required. The presence of the ITRs would confound such assemblies. Any structural variants covered by separate reads would be difficult to assemble, because it would be difficult to determine if any relevant variants occurring therein are cis or trans.

Since the AAV genome is approximately 4.8 kb in length [6], Pacific Biosciences (PacBio) sequencers are capable of accurately sequencing individual AAV molecules using their Circular Consensus algorithm [7]. Thus, it is possible to analyze any lot of manufactured AAV as a population of individuals without PCR or assembly bias.

Additional complications in the analysis of the sequencing data arise from the PacBio AAV library preparation protocol [11]. At the beginning of library preparation, the capsid needs to be lysed. After lysis, the single stranded viral DNA molecules must be manipulated into a double stranded form, which is a requirement prior to adapter ligation. For self complementary viral DNA, the molecule will form a snapback DNA with one double stranded end. For viral DNA that is not self complementary, incubating the DNA molecules to allow complementary molecules to anneal is included as a step in the protocol. But, it is possible for two viral DNAs with different sequences to anneal [12] provided that there is a sufficiently long region of complementary DNA. Thus, complete genomes and genome fragments can anneal as well as pairs of genomes that have differences. Additionally, the ITRs at the ends of the viral genome provide adequate complementarity to facilitate annealing. Even host cell DNA and other plasmid impurities can anneal with a viral genome if they share genetic elements.

The `--hd-finder` option in the PacBio CCS (Circular Consensus Sequence) program is normally used to find heteroduplexes, but this possibility can confound accurate counting of the viral DNA sequences.

Besides the annealing step, another complication arises from the gap filling step whereby a DNA polymerase is used. If there is extended incubation time in the protocol, the 3’ ITR end can serve as a primer for extending the single stranded viral genome into a double stranded DNA hairpin [13].

Also, the manufacturing process of AAVs can produce viruses containing partial genomes, DNA from the plasmids used in the manufacturing process, or DNA from the host cells [14].

Because of these complications, sequence analysis methods used for human genome resequencing do not apply for AAV. The alignment and variant calling algorithms will not properly account for structural variation or the variation in strandedness of the connected ITRs and payloads. In addition, the coverage found in AAV DNA sequencing is much larger than human genome resequencing and the variant frequency could be much smaller, even in the range of sequencing error.

In addition to the expected full-length vector genomes, the viral production process is known to generate a diverse array of rare and unexpected genome structures. To comprehensively capture this variability, we elected to analyze all DNA molecules obtained in a sequencing run, rather than focusing solely on the predominant species. This approach allows us to address the key question: “What are the components of each DNA molecule present in the sample, and how are they arranged?”. The tiling algorithm was developed specifically to answer this question.

Because our algorithm was not based on any structural presumptions, the algorithm can identify many complex structures. An examination of the infrequent tiling patterns at the tail of the example results deposited in Zenodo [19] reveal a remarkable diversity of genome structures. However, the results of the tiling algorithm require additional processing to organize and categorize them in order to provide insightful knowledge about the samples being sequenced. A recent preprint [20] describes such additional processing methods.

## Methods

The overarching principle of our methodology is to analyze each sequence individually and label them using a high-level description of the structure. These high-level descriptions are then grouped together, counted, and presented in descending order of abundance.

We use the verb, “tile”, in the one-dimensional abstraction of the tiling of a two dimensional surface. The tiles should cover the surface as completely as possible with a user-modifiable allowance of inter-tile gaps. That is, for a one-dimensional sequence, we want to maximally cover the sequence with subsequences derived from known sequences, see Fig 1, with minimal or no gaps.

**Fig 1.**
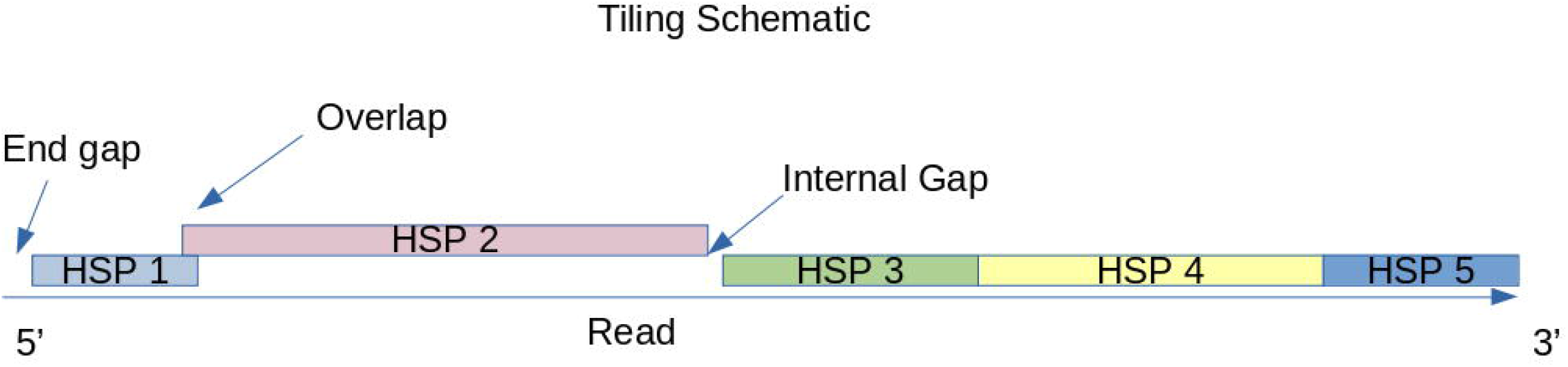
illustrates the tiling concept. The read is shown as a horizontal line from the 5’ to 3’ end, with colored blocks (HSP 1 to HSP 5) representing matches to known components. Ideally, these HSPs should fully span the read.

In the context of AAV DNA sequencing, it is also very important to understand the relationship between the “Flip” and “Flop” orientation of the ITR [15]. Specifically, if the 3’ ends of Flip and Flop are extended by complementing the base paired 5’ ends for 21 bases, that is the D region, Flip and Flop are revealed to be the exact reverse complement of each other. Thus, they can be treated as equivalent from a DNA sequence information perspective. We take advantage of this equivalence to simplify the analysis by using the single extended ITR sequence as a single Basic Local Alignment Search Tool (BLAST) [16] reference sequence.

Because the AAV replication process randomizes the orientation of the ITRs at each end of the viral genome combinatorially [10], conventional read alignment approaches, which rely on a fixed reference orientation, are not suitable. Instead, the tiling methodology uses the BLAST algorithm [16] to find regions of each read that match the expected components of a viral genome separately, and then uses this information to find coverage of the read with minimum gaps or overlaps.

Specifically, the reference BLAST database consists of the extended ITR sequence described above, the payload, and the backbone subsequences of the packaging plasmid for the viral genome. These subsequences in the reference BLAST database are referred to as ‘components’ in the succeeding text. Every read from the PacBio sequencing run is aligned against this reference and high scoring pairs (HSPs) of reference component and coordinates of the alignment are generated. All strandedness possibilities are included.

Thus, a more precise statement of the goal of the tiling algorithm is to find a set of HSPs for every read, sorted by their coordinates on the read, where the end of one HSP is juxtaposed against the start of the next HSP with no overlap or gap. In addition, the first HSP should have a starting read coordinate of 1, and the last HSP should have ending read coordinate equal to the length of the read.

Because the ITRs are self-complementing and many payloads are self-complementing or have repetitive sequences, many possible HSPs can be generated from the alignment of a read against the reference sequences. In addition, sequencing errors will complicate the analysis.

These issues are addressed in three steps. The first step is to assess if a tiled coverage is possible. This is done by creating a character string of blanks with the same length as the read. Then, “X”s are written into this string using string coordinates equal to the read coordinates of all the HSPs. If gaps of blank characters are greater in length than the `min_gap_search` parameter, then it is known that the read cannot be tiled successfully based on the reference components used. This coverage test can be modified by an optional set of reference sequences which will be described later. The default setting for the `min_gap_search` parameter is 11 bases. (N.B. the `min_gap_search` parameter is unrelated to the gap search in the BLAST algorithm.)

The second step is redundant HSP filtering. In this step, all the HSPs are sorted by the starting coordinates of the read and then by sorting on the ending coordinates of the read. Then, a search is made of any HSP that is fully contained within another HSP with respect to the read coordinates, regardless of the strandedness. There are two cases of containment to consider. The simplest case is where the containing HSP is larger than the contained HSP, in which case the contained HSP is then deleted from consideration.

The more complex case is when the two HSPs cover exactly the same coordinates of the read. In that case, the HSPs have different strandedness. This occurs frequently with ITRs due to their palindromic and repetitive composition. Here, we examine how many mismatches and gaps are found in each HSP and we pick the HSP that has the smallest sum of mismatches and gaps. If these sums are identical for both HSPs, then we just pick the HSP that uses the positive strand of the reference for the alignment so the process is deterministic.

The final step is to exhaustively search for a set of HSPs that have the best coverage score. In this context, “best” means the minimization of the following score, *S*

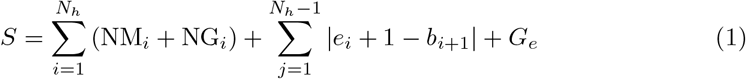

where *N*_*h*_ or is the number of HSPs, NM_*i*_ is the number of mismatches in HSP_*i*_, NG_*i*_ is the number of gaps in HSP_*i*_, *e*_*i*_ is the end coordinate of the read in HSP_*i*_, *b*_*i*_ is beginning coordinate of the read in HSP_*i*_ and *G*_*e*_ is the difference between the read length and the end coordinate of the read in the final HSP.

The first term in *S* captures the quality of the HSPs, and the second term captures the quality of the tiling coverage between HSPs. The third term captures any uncovered bases at the 3’ end. Any gap at the 5’ end of the read is constant because the HSPs are sorted by the beginning coordinate for the read in the HSP and overlapping HSPs are reduced to a single, largest possible HSP. Thus, any 5’ gap need not be considered in the minimization of the tiling score.

The exhaustive search is performed on a directed graph of HSPs where the root of the graph is the first HSP in the sorted order. Children of any node in the graph are those HSPs where the magnitude of the difference of parent node’s ending coordinate from the child node starting coordinate is less than the gap limit parameter. Fig 2 illustrates a simple, example graph.

**Fig 2.**
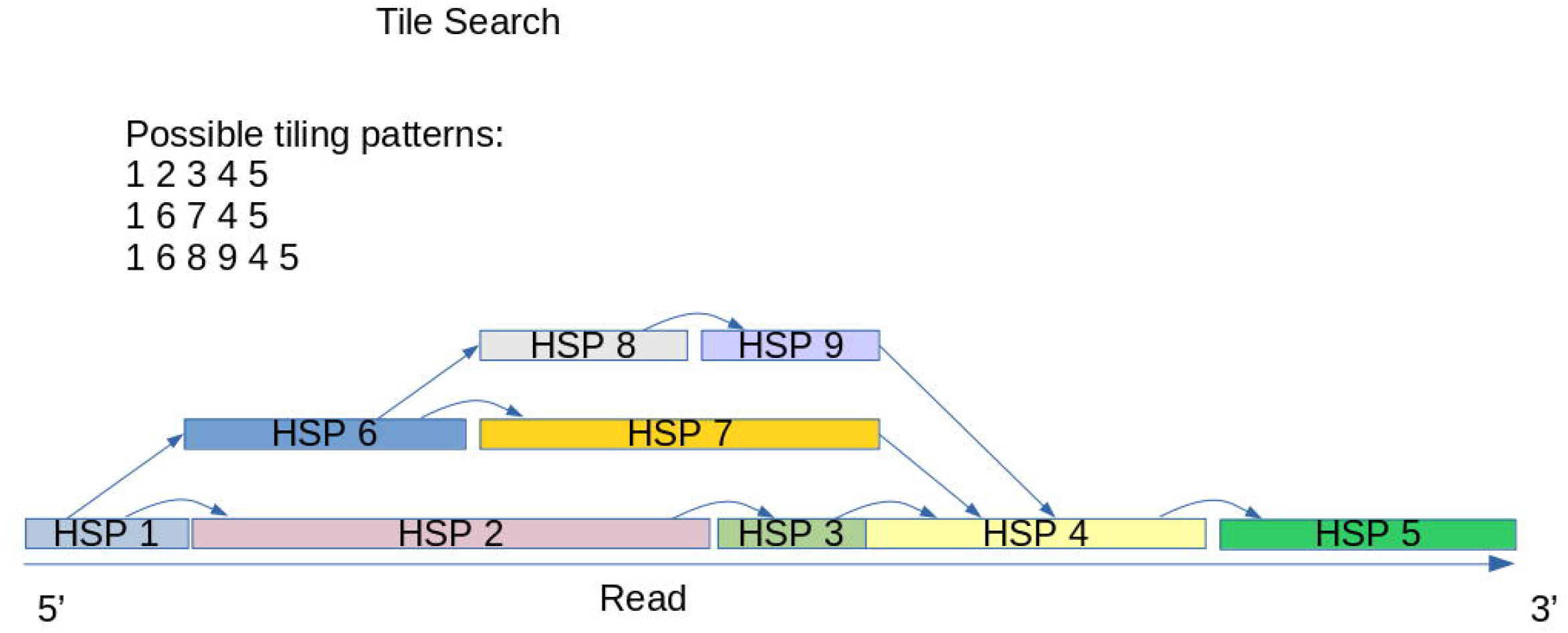
shows a simple graph of possible HSPs and the paths that can be taken to generate a complete tiling pattern.

If the HSP nodes in this graph have many linked HSPs, the exhaustive search can take a very long time. Thus, the algorithm counts the number of nodes visited in the search and aborts the search if the count exceeds the `max_iter` parameter. For good quality, less heterogeneous sequencing runs, the HSP search graph is usually very narrow, and can be searched very quickly.

Once the best coverage path through the HSPs is determined, then the code generates a summary description of the read by concatenating the BLAST database sequence name, the coordinates of the HSP match on the component along with the HSP strandedness. The summary description consists of a concatenation of all HSPs and which reference coordinates and strand each HSP aligns to. Note that unaligned bases in the original read are not represented in the summary description.

Here’s an example of a summary description of a read (from the 2021 dataset below, barcode bc1001):

~~~
ITR-FLIP[1-145](-) Payload[1-1831](-) ITR-FLIP[25-141](-) Payload[1-1831](+) ITR-FLIP[1-145](+)
~~~

From this example, one can see that this DNA sequence has self-complementary payload with complementary ITRs at the ends and a shortened ITR in the middle joining the double stranded payload. The coordinates shown in the tiling pattern reflect the coordinates of reference sequences. The strand character (+ or -) specify whether the matching sequence on the read is identical to the reference or is the reverse complement of the reference, respectively. The coordinates of these matches in the read are provided in the “.hsps” output file described below.

The code then counts the number of occurrences of each of these tiling patterns and sorts them by abundance in reverse order.

### Adjustments to the tiling process

There are some optional adjustments that can be made to the tiling algorithm’s process. First, the code can generate “pseudo” HSPs out of homopolymers. These “pseudo” HSPs are identified by applying a simple regular expression which looks for repeats of the same base. The minimum number of bases in the homopolymer can be set by the `min_poly` parameter. These HSPs have a component name of “poly-” followed by the base. Optionally, the length of the homopolymer can be added. These homopolymer HSPs are especially useful when poly-adenylation is used in a library preparation protocol.

Another adjustment is the use of multiple reference sequences after the initial reference database BLAST search. During the coverage checking step, if a large gap is found, the code will check other references for HSPs to see if the gaps can be covered. One such use case for additional reference sequences is the detection of additional plasmids used in AAV manufacturing. Occasionally, DNA from these plasmids will be incorporated into the capsid, and this additional reference check will identify them. In addition, contaminants such as host cell DNA and manufacturing plasmid DNA can also be incorporated into AAV DNA. By adding a human genome assembly and plasmid sequences as reference databases, HSPs with contaminant DNA can also be detected and included in the tiling patterns.

### Strand counting

A challenge with counting DNA molecules that is specific to AAV is the annealing protocol typically used to prepare sequencing libraries. During sequencing, a single SMRTbell molecule is sequenced multiple times by the same polymerase within a ZMW to generate multiple subreads that are combined to form a CCS read. When a SMRTbell molecule is generated from two fully complementary strands of the viral genomes (homoduplex), fewer polymerase passes (at least three passes) are sufficient to generate a high-confidence CCS read. However, a SMRTbell molecule generated from two different viral genomes with potential mismatches between complementary strands (heteroduplex molecules) increases sequencing error rates, and more polymerase passes (at least six) are required. Even with five passes, only one strand will be corrected, and the other may be missed. This potentially reduces the likelihood of generating a CCS read within the limited number of polymerase passes typically observed in a ZMW. Hence, heteroduplexes are less likely to meet CCS generation thresholds and will be under-counted relative to homoduplexes.

A second strand counting issue arises from the self-complementarity of the payload. After lysis of the capsid, a self complementary payload can anneal to itself forming a snapback DNA with one double stranded end. This end can be ligated to the PacBio SMRTBell adapter, thus generating a single, circularized DNA strand that can be sequenced. The PacBio Zero Mode Waveguide that binds this molecule will only sequence the DNA from one viral particle. In contrast, the expected common case for a single stranded AAV payload is that it anneals to another AAV viral genome, and thus, the Zero Mode Waveguide that sequences this pair of double stranded DNA molecules will be sequencing the DNA from two different viral particles.

In principal, it should be possible to discriminate between these two possibilities by using Unique Molecular Identifier (UMI) sequences attached to the SMRTbell adapter. The snapback DNA resulting a self-complementary genome would only have one UMI whereas the annealed genomes would have two.

Another artifact can arise from the 3’ end of the AAV ITR. When DNA polymerase is used in the library preparation for filling gaps, the 3’ end can prime the complete duplication of the AAV genome resulting in a large snapback DNA which can be ligated to a SMRTBell adapter. ZMW’s containing such snapback DNA’s represent a single AAV genome. These can be detected by the size of the CCS read, which will be approximately double the expected AAV genome.

We have not attempted to resolve this counting problem. Instead, the algorithm can generate two possible counts of tiling patterns. It will always generate a count based on the CCS reads. Optionally, it can generate a count based on what’s seen in each ZMW. If the ZMW reports just one read, then the output is just the tiling pattern for that read. However, if there are two reads reported, the tiling computes the pattern for both and reports them both.

## Implementation

The tiling algorithm is implemented in Perl using the Bio::Frescobi [17] Perl module which uses a relational database to store all the BLAST results and read sequences. The relational database greatly simplifies the retrieval of relevant HSPs for each read in an analysis. Frescobi supports either PostgreSQL or SQLite as the database engine. The code is available in Github [18].

An important property of the sequence database constructed by Frescobi is the 1:1 mapping of sequences to a short identifier, the `seqid`. Even if a sequence dataset has multiple read with identical sequences, each identical sequence will have the same identifier. Thus, any sequence based computation is only computed for unique sequences. Frescobi maintains a table to relate the name of a sequence to its identifier, so the abundance of sequences can be counted by finding all the names for a given seqid.

The script takes the name of sequence library in Frescobi as an input and can generate a large number of output files, see Table 1 for a list of commonly used output files. All of these files begin with the library name, and the extension indicates the type of data in the file. Certain execution options can generate additional files which are described with the tiling code documentation.

**Table 1.** Common Output files.

| Extension | Description of file contents |
| --- | --- |
| .tiles | Tile pattern for each unique read sequence. |
| .gap.seq | Subsequences in each read that were not identifiable. |
| .unmatched.seq | Full sequence of unique reads that had some sequence that was not identifiable. |
| .summary | Summary statistics of the tiling run. |
| .split.reads | Listing of sequences of each of the components for each unique read. |
| .tile.err | Any error messages. |
| .hsps | List of all HSPs for each unique read. |
| .tile.details | More details on each tiling pattern as a function of each unique read. |
| .name2seqid | The correspondence between `seqid` and the name of the sequence in the PacBio sequence file. This name identifies the ZMW number on the SMRT Cell. |
| .tile.zmw.counts | Tile patterns for the sequences per ZMW. |
| .feats | Counts for the features seen in the tiling run. |
| .tile.counts | Abundances for all the tile patterns sorted most abundant first. |
| .gap.counts | Abundances for the sequences that were not covered by reference sequences. This file is especially important in diagnosing what's unidentified in a sample. |

## Material

We analyzed four AAV datasets [Table 2] available on the PacBio web site, https://downloads.pacbcloud.com/public/dataset/AAV/ using the tiling algorithm. The files that we used have been archived in Zenodo [21].

**Table 2.** Listing of PacBio AAV datasets.

| Folder Name | Sample Name | Source |
| --- | --- | --- |
| 2021-scAAV-CBA-eGFP | 2021-scAAV/rh10-CBA-eGFP | Vector Biolabs |
| 2022-ssAAV-pAV-CMF-GFP | 2022-pAV-CMV-GFP | Vigene Biosciences |
| 2022-ssAAV-scAAV-mix | Same as folder | Not specified |
| 2023-Revio-ssAAV-pAV-CMF-GFP | Same as folder | Charles River |

## Results

The tiling algorithm was applied to the four AAV datasets, and the complete results were uploaded to Zenodo [19, 21]. In all cases, the number of unique tiling patterns was very large, and the analysis in this paper is limited to more abundant or notable patterns. Additional analysis tools that refine and aggregate the tiling algorithm patterns are demonstrated in Rouleau *et al* [20].

### 2021-scAAV-CBA-eGFP

This experiment had eight different samples that were multiplexed into one run. There is no information available on the differences between each sample. The percentage of sequences that could be characterized was very high across all eight samples with values between 99.4113% and 99.4849%.

We picked one sample arbitrarily for the display of detailed results. The sixteen most common patterns in the sample, barcoded bc1002, are given in Table 3

**Table 3.** Most abundant tiling patterns for 2021-scAAV-CBA-eGFP.

| Count | Frequency | Tiling pattern |
| --- | --- | --- |
| 41389 | 0.1288309 | ITR-FLIP[1-145](-) Payload[1-1831](-) ITR-FLIP[25-141](+) Payload[1-1831](+) ITR-FLIP[1-145](+) |
| 41325 | 0.1286317 | ITR-FLIP[1-145](-) Payload[1-1831](-) ITR-FLIP[25-141](-) Payload[1-1831](+) ITR-FLIP[1-145](+) |
| 40794 | 0.1269789 | ITR-FLIP[21-165](+) Payload[1-1831](-) ITR-FLIP[25-141](+) Payload[1-1831](+) ITR-FLIP[21-165](-) |
| 40623 | 0.1264466 | ITR-FLIP[21-165](+) Payload[1-1831](-) ITR-FLIP[25-141](-) Payload[1-1831](+) ITR-FLIP[21-165](-) |
| 18677 | 0.0581356 | ITR-FLIP[1-145](-) Payload[1-1831](-) ITR-FLIP[25-141](-) Payload[1-1831](+) ITR-FLIP[21-165](-) |
| 18535 | 0.0576936 | ITR-FLIP[1-145](-) Payload[1-1831](-) ITR-FLIP[25-141](+) Payload[1-1831](+) ITR-FLIP[21-165](-) |
| 17436 | 0.0542728 | ITR-FLIP[21-165](+) Payload[1-1831](-) ITR-FLIP[25-141](+) Payload[1-1831](+) ITR-FLIP[1-145](+) |
| 17362 | 0.0540424 | ITR-FLIP[21-165](+) Payload[1-1831](-) ITR-FLIP[25-141](-) Payload[1-1831](+) ITR-FLIP[1-145](+) |
| 5859 | 0.0182372 | ITR-FLIP[1-144](-) Payload[1-1831](-) ITR-FLIP[25-141](+) Payload[1-1831](+) ITR-FLIP[1-144](+) |
| 5816 | 0.0181034 | ITR-FLIP[1-144](-) Payload[1-1831](-) ITR-FLIP[25-141](-) Payload[1-1831](+) ITR-FLIP[1-144](+) |
| 5767 | 0.0179509 | ITR-FLIP[22-165](+) Payload[1-1831](-) ITR-FLIP[25-141](+) Payload[1-1831](+) ITR-FLIP[22-165](-) |
| 5458 | 0.0169890 | ITR-FLIP[22-165](+) Payload[1-1831](-) ITR-FLIP[25-141](-) Payload[1-1831](+) ITR-FLIP[22-165](-) |
| 1950 | 0.0060697 | ITR-FLIP[1-144](-) Payload[1-1831](-) ITR-FLIP[25-141](-) Payload[1-1831](+) ITR-FLIP[22-165](-) |
| 1884 | 0.0058643 | ITR-FLIP[1-144](-) Payload[1-1831](-) ITR-FLIP[25-141](+) Payload[1-1831](+) ITR-FLIP[22-165](-) |
| 1799 | 0.0055997 | ITR-FLIP[22-165](+) Payload[1-1831](-) ITR-FLIP[25-141](+) Payload[1-1831](+) ITR-FLIP[1-144](+) |
| 1731 | 0.0053881 | ITR-FLIP[22-165](+) Payload[1-1831](-) ITR-FLIP[25-141](-) Payload[1-1831](+) ITR-FLIP[1-144](+) |

The numbers on the first column represent how many times a pattern was tiled or the species existed in the sample. The second column represents the proportion of reads that have the particular pattern. As expected, we found majority of the patterns show a self complementary payload with a shortened ITR in the middle (no D region) and full length ITRs at the ends. The sequence differences between the patterns of similar abundance occur because of Flip or Flop ITR orientation or the strandedness of the shortened, interior ITR. The less abundant patterns show one missing base at the end of the ITR, 144 bases instead of 145. There are a total of 15,830 patterns seen, although many are just single occurrence patterns. The percentage of reads which had single occurrence patterns was 3.47%.

One expected observation in the results for 2021-scAAV-CBA-eGFP dataset was that the strandedness of the payload was constant. This is a consequence of the self complementary payload. The strandedness of the payload would be constant regardless of whether the library preparation operated on a snapback DNA with one double stranded end or two annealed, identical viral genomes.

### 2022-ssAAV-pAV-CMF-GFP

We were able to tile 93.7% of the reads from this dataset. There were a total of 173,677 patterns seen in this sample. 135,615 (78%) were single occurrence patterns, representing 20.79% of the sequence reads in the sample, and indicating a greater genome heterogeneity than the previous sample.

Shown in Table 4 is a partial listing of tiling patterns, in order of abundance, selected for interesting features:

**Table 4.**
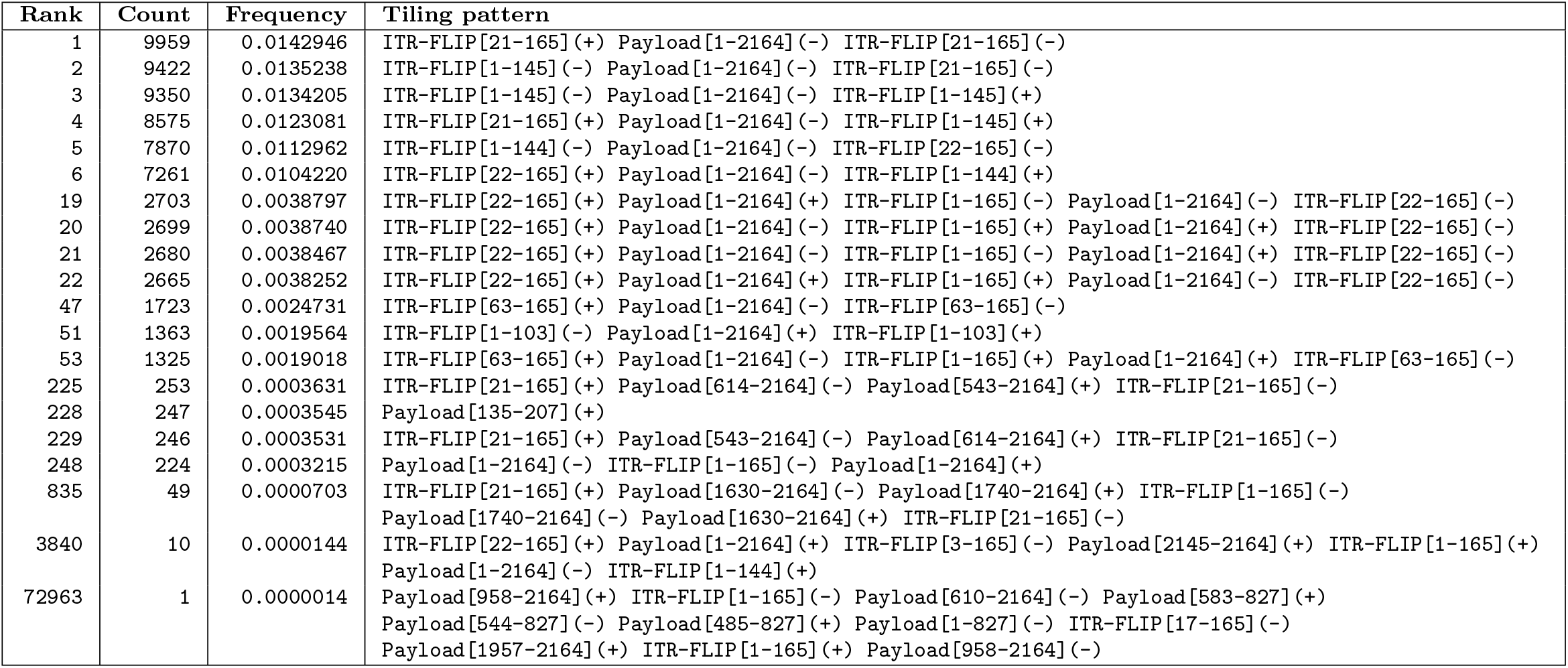
Selected tiling patterns for 2022-ssAAV-pAV-CMF-GFP.

The first four patterns are the most abundant. Unlike the previous sample where the most abundant pattern represented about 13% of the reads, the most abundant pattern in this sample represented only about 1.4% of the species present. The next two patterns (rank 5 and 6) show variation in the ITR’s: one missing base. In the 19^th^ rank, we observe the first of many patterns which indicate strand extension during library preparation. In the 47^th^ rank and lower, we see the first example of a large deletion in the ITRs. At rank 225, we find the first example of a “snapback” genome where the payload has folded back on itself to make a large hairpin. At rank 228, we find the first partial payload. Rank 229 is the inverse of rank 225, except for the ITRs. Rank 248 is the first complete payload with missing terminal ITRs. At rank 835, we see the first self complementary “snapback”, where the snapback sequence is replicated within these DNA molecules. Finally, an example of sequence that appears 10 times and one example of a sequence that appears once is included. Their patterns are far removed from the designed AAV genome.

This sample illustrated the great variety of structural patterns that can be discerned by the tiling algorithm. Given the large number of copies (except for the final example in Table 4), they are likely real. In addition, because we have the sequences of all these examples, PCR could be used to amplify these molecules for additional sequence verification.

The `README.txt` file accompanying these AAV datasets at Pacific Biosciences references the preprint by Talevich *et al* [23] for analysis. Only the 2021-scAAV-CBA-eGFP and 2022-ssAAV-pAV-CMF-GFP datasets were analyzed, and the results were found in the two Supplementary Files. Their method is based on using Minimap2 [24, 25] with references of the sequences for the expected AAV genome, RepCap and Helper plasmids, and host cell genome to generate primary alignments. Using these alignments, each read was assigned to a source with the ITR’s being separately analyzed using a Smith Waterman alignment to ITR sequences in both flip and flop orientation. In contrast, the tiling algorithm examines all the HSP’s between the reads and the references, and it searches for a combination of HSP’s that best covers each read, ideally with no gaps or overlaps.

The Talevich *et al* preprint reported the relative proportions of self-complementary and single stranded AAV genomes found in the two datasets. However, there is a discrepancy with both datasets with regard to the number of reads analyzed. For the 2021-scAAV-CBA-eGFP dataset, there are eight barcodes used. None of the eight sequence data files has a match of read number to the number of reads shown on page 4 of the their Supplementary File 2, which corresponds to the 2021-scAAV-CBA-eGFP dataset. Likewise for the 2022-ssAAV-pAV-CMF-GFP dataset, the number of reads shown on page 4 of the their Supplementary File 1 (877,543) did not match the number of reads for the reads in the sequence data file (652,367 reads).

Despite these discrepancies, we performed an analysis using the tiling results to see if we obtained comparable values for the single stranded and self-complementary patterns. We binned the tiling results for these two datasets into three categories; self-complementary AAV, single stranded AAV, and other. The categorization was performed by the script, `summarize sc_vs_ss.pl`, which is included in the Zenodo archive [21]. The binning was performed by simplifying the tiling patterns into just ITR’s and payloads. If it finds the pattern of “ITR Payload ITR”, it is counted as “single stranded”. If it finds a pattern of “ITR Payload ITR Payload ITR” where the two payload components are opposite strandedness, it counted as “self complementary”. All other patterns are counted as “other”. For the 2021-scAAV-CBA-eGFP dataset, we analyzed sequences with all eight barcodes. A comparison of the results in shown in Table 5.

**Table 5.** Comparison of AAV types between Talevich *et al* and tiling.

| **Dataset:** | **2021-scAAV-CBA-eGFP** |  |  | **2022-ssAAV-pAV-CMF-GFP** |  |  |
| --- | --- | --- | --- | --- | --- | --- |
| **Method** | **Self Comp.** | **Single Strand** | **SC/Sum Ratio** | **Self Comp.** | **Single Strand** | **SC/Sum Ratio** |
| Talevich *et al* | 258069 | 4449 | 0.983 | 314466 | 512382 | 0.380 |
| Tiling | 2262398 | 12686 | 0.994 | 163711 | 250288 | 0.395 |

As can be seen in Table 5, the ratios of self-complementary to the sum of both AAV bins are similar.

### 2022-ssAAV-scAAV-mix

This run was apparently a mixture test using the two samples above. The reference sequence used for tiling was ITR-FLIP plus the payload and backbones for the above two viral genomes. The percentage of reads that could be tiled was high: 98.949%.

The 24 most abundant tiling patterns are listed in Table 6.

**Table 6.** Selected tiling patterns for 2022-ssAAV-scAAV-mix.

| Count | Frequency | Tiling pattern |
| --- | --- | --- |
| 27416 | 0.0910731 | ITR-FLIP[1-145](-) Payload_scAAV[1-1831](-) ITR-FLIP[25-141](+) Payload_scAAV[1-1831](+) ITR-FLIP[1-145](+) |
| 27051 | 0.0898606 | ITR-FLIP[1-145](-) Payload_scAAV[1-1831](-) ITR-FLIP[25-141](-) Payload_scAAV[1-1831](+) ITR-FLIP[1-145](+) |
| 26806 | 0.0890467 | ITR-FLIP[21-165](+) Payload_scAAV[1-1831](-) ITR-FLIP[25-141](+) Payload_scAAV[1-1831](+) ITR-FLIP[21-165](-) |
| 26598 | 0.0883558 | ITR-FLIP[21-165](+) Payload_scAAV[1-1831](-) ITR-FLIP[25-141](-) Payload_scAAV[1-1831](+) ITR-FLIP[21-165](-) |
| 14035 | 0.0466228 | ITR-FLIP[1-145](-) Payload_scAAV[1-1831](-) ITR-FLIP[25-141](-) Payload_scAAV[1-1831](+) ITR-FLIP[21-165](-) |
| 13948 | 0.0463338 | ITR-FLIP[1-145](-) Payload_scAAV[1-1831](-) ITR-FLIP[25-141](+) Payload_scAAV[1-1831](+) ITR-FLIP[21-165](-) |
| 13371 | 0.0444171 | ITR-FLIP[21-165](+) Payload_scAAV[1-1831](-) ITR-FLIP[25-141](+) Payload_scAAV[1-1831](+) ITR-FLIP[1-145](+) |
| 13362 | 0.0443872 | ITR-FLIP[21-165](+) Payload_scAAV[1-1831](-) ITR-FLIP[25-141](-) Payload_scAAV[1-1831](+) ITR-FLIP[1-145](+) |
| 5493 | 0.0182472 | ITR-FLIP[1-148](-) Payload_scAAV[1-1831](-) ITR-FLIP[25-141](+) Payload_scAAV[1-1831](+) ITR-FLIP[1-148](+) |
| 5265 | 0.0174898 | ITR-FLIP[1-148](-) Payload_scAAV[1-1831](-) ITR-FLIP[25-141](-) Payload_scAAV[1-1831](+) ITR-FLIP[1-148](+) |
| 5122 | 0.0170147 | ITR-FLIP[18-165](+) Payload_scAAV[1-1831](-) ITR-FLIP[25-141](+) Payload_scAAV[1-1831](+) ITR-FLIP[18-165](-) |
| 5058 | 0.0168021 | ITR-FLIP[18-165](+) Payload_scAAV[1-1831](-) ITR-FLIP[25-141](-) Payload_scAAV[1-1831](+) ITR-FLIP[18-165](-) |
| 2555 | 0.0084874 | ITR-FLIP[1-148](-) Payload_scAAV[1-1831](-) ITR-FLIP[25-141](+) Payload_scAAV[1-1831](+) ITR-FLIP[18-165](-) |
| 2490 | 0.0082715 | ITR-FLIP[1-148](-) Payload_scAAV[1-1831](-) ITR-FLIP[25-141](-) Payload_scAAV[1-1831](+) ITR-FLIP[18-165](-) |
| 2428 | 0.0080656 | ITR-FLIP[18-165](+) Payload_scAAV[1-1831](-) ITR-FLIP[25-141](+) Payload_scAAV[1-1831](+) ITR-FLIP[1-148](+) |
| 2377 | 0.0078961 | ITR-FLIP[18-165](+) Payload_scAAV[1-1831](-) ITR-FLIP[25-141](-) Payload_scAAV[1-1831](+) ITR-FLIP[1-148](+) |
| 1802 | 0.0059861 | ITR-FLIP[1-145](-) Payload_pAV_CMV_GFP[1-2164](+) ITR-FLIP[1-145](+) |
| 1790 | 0.0059462 | ITR-FLIP[21-165](+) Payload_pAV_CMV_GFP[1-2164](+) ITR-FLIP[21-165](-) |
| 1748 | 0.0058067 | ITR-FLIP[21-165](+) Payload_pAV_CMV_GFP[1-2164](-) ITR-FLIP[21-165](-) |
| 1731 | 0.0057502 | ITR-FLIP[21-165](+) Payload_pAV_CMV_GFP[1-2164](+) ITR-FLIP[1-145](+) |
| 1698 | 0.0056406 | ITR-FLIP[1-145](-) Payload_pAV_CMV_GFP[1-2164](+) ITR-FLIP[21-165](-) |
| 1697 | 0.0056373 | ITR-FLIP[1-145](-) Payload_pAV_CMV_GFP[1-2164](-) ITR-FLIP[1-145](+) |
| 1696 | 0.0056339 | ITR-FLIP[1-145](-) Payload_pAV_CMV_GFP[1-2164](-) ITR-FLIP[21-165](-) |
| 1685 | 0.0055974 | ITR-FLIP[21-165](+) Payload_pAV_CMV_GFP[1-2164](-) ITR-FLIP[1-145](+) |

The tiling patterns were analyzed by a short Perl script which counted the number of reads that had either one of the two AAV payloads, both, or neither. The self complementary AAV percentage was 85.814% and the single stranded percentage was 14.061%, with the ratio being 6.103. A small fraction (0.074%) were ITR fragments, and another small fraction (0.052%) had both sequences from both libraries, although the maximum number of copies seen for this category was 2 copies out of 301,033 total reads. The percentage of reads which had single occurrence patterns was 6.96%.

For experiments where assessment of mixture ratios for different AAVs is important, the tiling algorithm could be very useful, because each read is characterized independently. As long as each AAV has unique sequences relative to the other AAVs in the mixture, the ratio of counts could be definitive.

However, the strand counting issue, described earlier, could be a serious problem if the two AAV’s are different with respect to the self complementarity of the payload, that is if one is single stranded and one is self complementary, which is the case for 2022-ssAAV-scAAV-mix.

Since the `README.txt` file in the PacBio web site stated that the mixture ratio should have been 10 parts self-complementary library to 1 part single stranded, the measurement above was a discrepancy. The counting problem described above would exacerbate the discrepancy. If we assume that all single stranded genomes are annealed before binding into a ZMW, the counts for the single stranded genomes should be doubled. In contrast, if we assume that the self-complementary genomes will fold on themselves, then the counts for the self-complementary genomes should not be adjusted. If we make the above assumptions about the annealing of single stranded and self-complementary genome, then the ratio is 3.052.

### 2023-Revio-ssAAV-pAV-CMF-GFP

Two replicates were sequenced, but there’s no indication that they are different.

The percentage of sequences that could be tiled was lower than the previous three datasets with replicate 1 being 89.054% and replicate 2 being 87.4374%. The number of unique tiling patterns for each replicate are 904,153 for replicate 1 and 1,488,692 for replicate 2. The percentage of reads which had single occurrence patterns was 58.22%.

The most abundant 24 tile patterns for the two replicates are listed in tables 7 and 8. Compared to the previous samples, these sequencing runs showed much more heterogeneity in the tiling results, as indicated by the number of unique tiling patterns. More than half of the reads have unique tiling patterns. In addition, the most abundant patterns have a smaller frequency of occurrence relative to the tiling patterns for the other datasets (e.g. 0.9% for the first replicate versus 12.9% in the 2021-scAAV-CBA-eGFP for the most abundant pattern in the sample with the bc1002 barcode). We also see examples of strand extension on the ITRs in the last tiling patterns in tables 7 and 8.

**Table 7.** Replicate 1 most abundant tiling patterns for 2023-Revio-ssAAV-pAV-CMF-GFP.

| Count | Frequency | Tiling pattern |
| --- | --- | --- |
| 14561 | 0.0089000 | ITR-FLIP[21-165](+) Payload[1-2164](-) ITR-FLIP[21-165](-) |
| 14496 | 0.0088603 | ITR-FLIP[1-145](-) Payload[1-2164](-) ITR-FLIP[1-145](+) |
| 11057 | 0.0067583 | ITR-FLIP[22-165](+) Payload[1-2164](-) ITR-FLIP[22-165](-) |
| 10774 | 0.0065853 | ITR-FLIP[1-144](-) Payload[1-2164](-) ITR-FLIP[22-165](-) |
| 10729 | 0.0065578 | ITR-FLIP[21-165](+) Payload[1-2164](+) ITR-FLIP[21-165](-) |
| 10707 | 0.0065444 | ITR-FLIP[1-144](-) Payload[1-2164](-) ITR-FLIP[1-144](+) |
| 10667 | 0.0065199 | ITR-FLIP[1-145](-) Payload[1-2164](+) ITR-FLIP[1-145](+) |
| 10513 | 0.0064258 | ITR-FLIP[1-145](-) Payload[1-2164](-) ITR-FLIP[21-165](-) |
| 9666 | 0.0059081 | ITR-FLIP[22-165](+) Payload[1-2164](-) ITR-FLIP[1-144](+) |
| 9598 | 0.0058665 | ITR-FLIP[21-165](+) Payload[1-2164](-) ITR-FLIP[1-145](+) |
| 8350 | 0.0051037 | ITR-FLIP[1-144](-) Payload[1-2164](+) ITR-FLIP[22-165](-) |
| 7987 | 0.0048818 | ITR-FLIP[1-145](-) Payload[1-2164](+) ITR-FLIP[21-165](-) |
| 7821 | 0.0047804 | ITR-FLIP[1-144](-) Payload[1-2164](+) ITR-FLIP[1-144](+) |
| 7761 | 0.0047437 | ITR-FLIP[22-165](+) Payload[1-2164](+) ITR-FLIP[22-165](-) |
| 7621 | 0.0046581 | ITR-FLIP[21-165](+) Payload[1-2164](+) ITR-FLIP[1-145](+) |
| 7560 | 0.0046208 | ITR-FLIP[22-165](+) Payload[1-2164](+) ITR-FLIP[1-144](+) |
| 2194 | 0.0013410 | ITR-FLIP[1-144](-) Payload[1-2164](-) ITR-FLIP[21-165](-) |
| 2142 | 0.0013092 | ITR-FLIP[1-144](-) Payload[1-2164](-) ITR-FLIP[1-145](+) |
| 2036 | 0.0012445 | ITR-FLIP[22-165](+) Payload[1-2164](-) ITR-FLIP[21-165](-) |
| 2026 | 0.0012383 | ITR-FLIP[22-165](+) Payload[1-2164](-) ITR-FLIP[1-165](-) Payload[1-2164](+) ITR-FLIP[22-165](-) |
| 1983 | 0.0012121 | ITR-FLIP[1-144](-) Payload[1-2164](+) ITR-FLIP[1-165](-) Payload[1-2164](-) ITR-FLIP[1-144](+) |
| 1958 | 0.0011968 | ITR-FLIP[1-144](-) Payload[1-2164](-) ITR-FLIP[1-165](-) Payload[1-2164](+) ITR-FLIP[1-144](+) |
| 1942 | 0.0011870 | ITR-FLIP[1-144](-) Payload[1-2164](-) ITR-FLIP[1-165](+) Payload[1-2164](+) ITR-FLIP[1-144](+) |
| 1910 | 0.0011674 | ITR-FLIP[22-165](+) Payload[1-2164](-) ITR-FLIP[1-165](+) Payload[1-2164](+) ITR-FLIP[22-165](-) |

**Table 8.** Replicate 2 most abundant tiling patterns for 2023-Revio-ssAAV-pAV-CMF-GFP.

| Count | Frequency | Tiling pattern |
| --- | --- | --- |
| 17604 | 0.0064796 | ITR-FLIP[21-165](+) Payload[1-2164](-) ITR-FLIP[21-165](-) |
| 17030 | 0.0062683 | ITR-FLIP[1-145](-) Payload[1-2164](-) ITR-FLIP[1-145](+) |
| 13788 | 0.0050750 | ITR-FLIP[21-165](+) Payload[1-2164](+) ITR-FLIP[21-165](-) |
| 13523 | 0.0049775 | ITR-FLIP[1-145](-) Payload[1-2164](+) ITR-FLIP[1-145](+) |
| 12799 | 0.0047110 | ITR-FLIP[22-165](+) Payload[1-2164](-) ITR-FLIP[22-165](-) |
| 12770 | 0.0047003 | ITR-FLIP[1-144](-) Payload[1-2164](-) ITR-FLIP[22-165](-) |
| 12383 | 0.0045579 | ITR-FLIP[1-144](-) Payload[1-2164](-) ITR-FLIP[1-144](+) |
| 12366 | 0.0045516 | ITR-FLIP[1-145](-) Payload[1-2164](-) ITR-FLIP[21-165](-) |
| 11335 | 0.0041721 | ITR-FLIP[21-165](+) Payload[1-2164](-) ITR-FLIP[1-145](+) |
| 11173 | 0.0041125 | ITR-FLIP[22-165](+) Payload[1-2164](-) ITR-FLIP[1-144](+) |
| 10391 | 0.0038247 | ITR-FLIP[1-145](-) Payload[1-2164](+) ITR-FLIP[21-165](-) |
| 10341 | 0.0038063 | ITR-FLIP[1-144](-) Payload[1-2164](+) ITR-FLIP[22-165](-) |
| 9959 | 0.0036657 | ITR-FLIP[22-165](+) Payload[1-2164](+) ITR-FLIP[22-165](-) |
| 9802 | 0.0036079 | ITR-FLIP[1-144](-) Payload[1-2164](+) ITR-FLIP[1-144](+) |
| 9597 | 0.0035324 | ITR-FLIP[22-165](+) Payload[1-2164](+) ITR-FLIP[1-144](+) |
| 9549 | 0.0035147 | ITR-FLIP[21-165](+) Payload[1-2164](+) ITR-FLIP[1-145](+) |
| 3781 | 0.0013917 | ITR-FLIP[22-165](+) Payload[1-2164](-) ITR-FLIP[1-165](-) Payload[1-2164](+) ITR-FLIP[22-165](-) |
| 3731 | 0.0013733 | ITR-FLIP[1-144](-) Payload[1-2164](+) ITR-FLIP[1-165](-) Payload[1-2164](-) ITR-FLIP[1-144](+) |
| 3720 | 0.0013692 | ITR-FLIP[22-165](+) Payload[1-2164](-) ITR-FLIP[1-165](+) Payload[1-2164](+) ITR-FLIP[22-165](-) |
| 3711 | 0.0013659 | ITR-FLIP[1-144](-) Payload[1-2164](-) ITR-FLIP[1-165](+) Payload[1-2164](+) ITR-FLIP[1-144](+) |
| 3677 | 0.0013534 | ITR-FLIP[1-144](-) Payload[1-2164](-) ITR-FLIP[1-165](-) Payload[1-2164](+) ITR-FLIP[1-144](+) |
| 3598 | 0.0013243 | ITR-FLIP[22-165](+) Payload[1-2164](+) ITR-FLIP[1-165](-) Payload[1-2164](-) ITR-FLIP[22-165](-) |
| 3544 | 0.0013045 | ITR-FLIP[1-144](-) Payload[1-2164](+) ITR-FLIP[1-165](+) Payload[1-2164](-) ITR-FLIP[1-144](+) |
| 3437 | 0.0012651 | ITR-FLIP[22-165](+) Payload[1-2164](+) ITR-FLIP[1-165](+) Payload[1-2164](-) ITR-FLIP[22-165](-) |

The analysis of these AAV DNA sequences in these sequencing libraries using the tiling algorithm revealed a very high level of diversity in the population of viral DNAs.

Many of them are low abundance, but not single occurrence. Here’s an example of the most abundant snapback DNA from the second replicate of the 2023 Revio run:

~~~
1460 0.0005374 ITR-FLIP[21-165](+) Payload[1740-2164](-) Payload[1630-2164](+) ITR-FLIP[21-165](-)
~~~

The fact that there were 1460 copies of this DNA structure strongly suggests that it is not an computational artifact.

The percentage of reads that could not be tiled was greater than 10% in both replicates. Because the tiling algorithm provides frequency of occurrence measurements for the uncovered portions of the reads, an exploration of these sequences was conducted. In the uncompressed version of the file, `tiling/2023-Revio-ssAAV-pAV-CMF-GFP/phase1/rep1.gap.counts.gz` which is in the Zenodo archive [19], we found the following sequence,

AGGAACCCCTAGTGATGGAGTTGGCCACTCCCTCTCTGCGCGCTCGCTCG CTCACTGAGGCCGGGCGACCAAAGGTCGCCCGACGCCCGGGCTTTGCCCG GGCGTCGGGCGACCTTTGGTCGCCCGGCCTCAGTGAGCGAGCGAGCGCGC AGAGAGGGAGTGGCCAACTCCATCACTAGGGGTTCCT

This sequence has many common subsequences with the ITR sequence, but it is predicted by the RNAfold algorithm in the ViennaRNA Web Services [26, 27] to be snapback DNA with only the middle TTT bases forming a loop. We labeled this sequence, “BITR”, mnemonic for “Big ITR”. An additional ITR like sequence was also found with the same long hairpin pattern, but not included as a new reference.

Another sequence that was not found in the reference genome which appeared 450 times in replicate 1 is given below:

CAGAAGAACTCGTCAAGAAGGCGATAGAAGGCGATGCGCTGCGAATCGGGAGCGGCGATACCGTAAAGCACGAGGAAGCGGTCAGCCCATTCGCCGCCAAGCTCTTCAGCAATATCACGGGTAGCCAACGCTATGTCCTGATAGCGGTCCGCCACACCCAGCCGGCCACAGTCGATGAATCCAGAAAAGCGGCCATTTTCCACCATGATATTCGGCAAGCAGGCATCGCCATGGGTCACGACGAGATCCTCGCCGTCGGGCATGCTCGCCTTGAGCCTGGCGAACAGTTCGGCTGGCGCGAGCCCCTGATGCTCTTCGTCCAGATCATCCTGATCGACAAGACCGGCTTCCATCCGAGTACGTGCTCTCTCGATGCGATGTTTCGCTTGGTGGTCGAATGGGCAGGTAGCCGGATCAAGCGTATGCAGCCGCCGCATTGCATCAGCCATGATGGATACTTTCTCGGCAGGAGCAAGGTGAGATGACAGGAGATCCTGCCCCGGCACTTCGCCCAATAGCAGCCAGTCCCTTCCCGCTTCAGTGACAACGTCGAGTACAGCTGCGCAAGGAACGCCCGTCGTGGCCAGCCACGATAGCCGCGCTGCCTCGTCTTGCAGTTCATTCAGGGCACCGGACAGGTCGGTCTTGACAAAAAGAACCGGGCGCCCCTGCGCTGACAGCCGGAACACG

This sequence was selected by examining the `rep1.gap.counts` file which contains the sequences that did not match the reference genome. This file is sorted in reverse order of abundance. This sequence was much longer than any others that were more abundant. We gave this sequence the name of “S450” simply based on the occurrence count.

We performed several Blast searches at NCBI using the ‘Blastn’ algorithm against the “Core NT” database and found MZ504910.1 [28] as a good hit. We then used the ‘Blastn’ algorithm against the Patented sequence using the Big ITR sequence above and found OF452498.1 [29] which was a patented AAV sequence. We also downloaded the helper cloning vector, AF369965.1, [30] as a likely source of incorporated vector genomes.

Using these four additional sequence files as references, we then reran both Blastn and the tiling algorithm where these four references were used in addition. The updated tiling results are included in Zenodo [21]. The most frequent tiling patterns were reproduced with the addition of the four references, and the percentage of the tiled reads improved. In replicate 1, the percentage increased from 89.054% to 92.622%, and in replicate 2, from 87.437% to 91.3037%. The percentage of reads with single occurrence patterns remained substantial (57.54%)

The presence of these additional references in the tiling patterns for this AAV sample are shown in Table 9

**Table 9.** Presence of Other References in Reads.

| Reference ID | Number of Reads | Percentage of Tiled Reads |
| --- | --- | --- |
| BITR | 10890 | 0.27 |
| pHelper | 23217 | 0.58 |
| MZ504910.1 | 17325 | 0.43 |
| OF452498.1 | 351402 | 8.79 |

It should be noted that these additional references were selected non-systematically. The intention was to illustrate how the unidentified sequences could be used to further characterize this AAV sample.

Because we do not know the details of how this sample was produced, our analysis of the unidentified sequences is quite incomplete. Ideally, the sequences of the plasmids and cell lines used by the producer of this sample would be used as additional references to help characterize the unidentified sequences.

## Discussion

A key strength of our approach is its ability to identify noncanonical and complex genome arrangements by exhaustively parsing tiling patterns using high-scoring pairs (HSPs) from BLAST alignment against annotated reference components. This allows us to map every segment within each CCS read including strand orientation and position to reconstruct a comprehensive genome layout. By segregating repetitive elements in the reference BLAST database, we can properly annotate repetitive elements in a small genome.

Alternative AAV genome classification frameworks were developed to address similar challenges [22, 23]. The pipelines typically combine HiFi CCS read generation with alignment via Minimap2 and use BLAST-based parsing in Python to classify sequences into structural categories such as full genomes, snapback genomes, genome deletion mutants, and other noncanonical forms.

In contrast, our pipeline provides a more granular, tile-based reconstruction of each AAV genome by aligning and identifying full-length reference components including ITRs, payloads, and backbones using BLAST. Rather than relying solely on coordinate overlaps or a primary alignment, our method maps all segments within each read to reconstruct the complete genomic configuration. This includes accurate determination of components, their count, strand orientation, and completeness (full or partial).

For instance, while other methods may flag an ITR based on a short alignment, our approach ensures that only biologically meaningful and structurally complete ITRs are recognized. Furthermore, by reporting all unique tiling patterns and their abundances, our pipeline enables high-resolution insight into both dominant and low-frequency structural variants, offering a better view of the diversity within AAV vector preparations and reducing the data complexity by approximately two orders of magnitude.

In addition to identifying well-matched patterns, the tiling algorithm also reports reads or parts of reads that don’t match anything in the reference. These “untiled” reads and unmatched parts are saved in output files (like.gap.seq,.gap.counts, and.unmatched.seq), allowing scientists to look more closely at unexpected or unknown sequences that may come from contamination or missing reference information. This capability was illustrated in the additional analysis of the 2023-Revio-ssAAV-pAV-CMF-GFP sample.

The use of an exhaustive search to find the best tiling pattern is potentially a problem for the general use of the tiling algorithm. In particular, repetitive elements in the reference sequence will result in many overlapping HSPs, and the search will not complete in a reasonable time.

The challenge with the tiling analysis is the high level of detail. Typical tiling runs can generate tens of thousands of patterns, but many of them are from very low abundance DNAs (literally parts per million). In addition, the coordinates in the HSPs are affected by sequencing errors, and small differences in the endpoints are typically not significant. Thus, there is great value in processing the tiling output to summarize the patterns that are seen. Rouleau et. al. [20] describe analytic methods that provide additional insights into the composition of a sequenced viral genome sample.

Although demonstrated here in the context of adeno-associated virus (AAV) genome analysis, the tiling algorithm is generalizable and can be applied to other DNA molecules with known component structures. For example, many viral genomes and eukaryotic gene transcripts can be sequenced as high fidelity long reads, and thus can be analyzed using the tiling algorithm. Nonetheless, certain design features may limit its broader applicability as discussed above. It is important to note that the algorithm does not perform base-pair level sequence analysis or variant calling. Instead, it provides high-level structural annotations of each DNA molecule, identifying the order, orientation, and boundaries of predefined genomic components such as inverted terminal repeats (ITRs), payloads, plasmid backbone elements, etc. These structural annotations form a framework for downstream analyses. When nucleotide-level resolution is required, the algorithm’s output coordinates can be used to extract relevant subsequences from individual reads. The software implementation includes functionality to generate these coordinates and subsequences for each annotated molecule.

## Conclusion

We have presented an algorithm for structural analysis of DNA sequences that can handle rearrangements on individual AAV molecules. The algorithm also provides the mechanism to identify unexpected components in a DNA sample as well as provide some quantification of proportions in a mixture. Designed originally for AAV genome analysis, our algorithm can be utilized with any DNA molecules with known component substructures and structural variability.

## Acknowledgments

We thank James McGivney for his leadership in supporting AAV sequencing at Oxford Biomedica. We thank Jeong-Ah Kwon, Brianna Uga, Esra D. Camci, and Conner Traugot for their laboratory work in preparing samples and running the Sequel II instrument to provide high quality sequence data for analysis.

We thank Pacific Biosciences for technical assistance with processing sequence data and smooth functioning of our Sequel II instrument as well as permission to archive the AAV dataset in Zenodo.

We thank the reviewers and editor at PLOS ONE for their insightful reviews. Addressing their comments has materially improved the quality of our paper.

